# Phylogeny, ancestors and anagenesis in the hominin fossil record

**DOI:** 10.1101/434894

**Authors:** Caroline Parins-Fukuchi, Elliot Greiner, Laura M. MacLatchy, Daniel C. Fisher

## Abstract

Probabilistic approaches to phylogenetic inference have recently gained traction in paleontological studies. Because they directly model processes of evolutionary change, probabilistic methods facilitate a deeper assessment of variability in evolutionary pattern by weighing evidence for competing models. Although phylogenetic methods used in paleontological studies have generally assumed that evolution proceeds by splitting cladogenesis, extensions to previous models help explore the potential for morphological and temporal data to provide differential support for contrasting modes of evolutionary divergence. Recent methodological developments have integrated ancestral relationships into probabilistic phylogenetic methods. These new approaches rely on parameter-rich models and sophisticated inferential methods, potentially obscuring the respective contributions of data and models. In this study, we describe a simple likelihoodist approach that combines probabilistic models of morphological evolution and fossil preservation to reconstruct both cladogenetic and anagenetic relationships. By applying this approach to a dataset of fossil hominins, we demonstrate the capability of existing models to unveil evidence for anagenesis presented by morphological and temporal data. This evidence was previously recognized by qualitative assessments, but largely ignored by quantitative phylogenetic analyses. For example, we find support for directly ancestral relationships in multiple lineages: *Sahelanthropus* is ancestral to later hominins; *Australopithecus anamensis* is ancestral to *Au. afarensis*; *Au. garhi* is ancestral to *Homo*; *H. antecessor* is ancestral to *H. heidelbergensis*, which in turn is ancestral to both *H. sapiens* and *H. neanderthalensis*. These results show a benefit of accommodating direct ancestry in phylogenetics. By so doing, quantitative results align more closely with previous qualitative expectations.

## Introduction

Phylogenetic methods that rely upon probabilistic evolutionary models have recently begun to emerge as important tools for addressing paleobiological issues at macroevolutionary timescales (Wright and Hillis 2014; Puttick et al. 2017). Probabilistic approaches appear promising in their performance relative to earlier cladistic methods and offer several important advantages. These include their tendency to increase the clarity with which models are specified and their ability to weigh evidence under competing evolutionary models using modern inferential machinery. While probabilistic approaches have long been dominant in molecular phylogenetics, their application to paleontological datasets is largely untapped. Yet the extension of approaches developed for extant taxa to fossil taxa will allow for the testing of a broader range of evolutionary hypotheses that are designed to reflect differences in the morphological patterns underlying divergence between fossil taxa.

Most previous approaches to phylogenetic reconstruction depict evolutionary relatedness using patterns of ‘hypothetical’ ancestry in which more closely related taxa are represented as possessing more recent common (hypothetical) ancestors. However, when fossil taxa are included in phylogenetic analyses, we might reasonably consider whether direct ancestry between temporally disjunct taxa or samples can be detected (Foote 1996). Although there is a spectrum of possible patterns of temporal distribution reflecting directly ancestral relationships, these may be dichotomized into cladogenesis and anagenesis.

Characterization of the relative prevalence of cladogenesis and anagenesis in the fossil record and their bearing on the study of evolutionary mode has long been a fundamental question in paleobiology (Simpson 1944; Stanley 1998; Levinton 2001). Although the significance of these contrasting models of evolutionary change in interpreting patterns in the fossil record has been fiercely debated (Gould and Eldredge 1977; Gould 1980; Levinton and Chris 1980; Gingerich 1985; Futuyma 1987), anagenetic change has been widely observed in the fossil record and can impact inferences of evolutionary processes (MacLeod 1991). This may be particularly important when constructing and testing hypotheses of phylogenetic relationships, where accommodation of anagenetic change may improve accuracy and yield insights otherwise unattainable (Gingerich 1979; Fox et al. 1999; Aze et al. 2011; Strotz and Allen 2013; Aze et al. 2013).

Quantitative phylogenetic approaches have usually assumed the universality of cladogenesis. Until recently, this fundamental limitation has remained largely intact, despite the development of increasingly sophisticated probabilistic approaches to phylogenetic inference. Since most of the major developments in probabilistic approaches to phylogenetics were developed for molecular data (but see Lewis 2001), their use has been largely restricted to neontological applications. As a result, there has been relatively little impetus in the areas most engaged in phylogenetic research to consider phyletic evolution, leaving the role of anagenesis in phylogenetic reconstruction under-explored.

The conceptual issues discussed above are particularly relevant in taxa such as the hominin clade (humans and all other taxa more closely related to humans than to genus *Pan*), where hypotheses of direct ancestry are often proposed. Discerning human evolutionary patterns is a focus traceable to Darwin (1871), and ancestry is almost certainly more frequently proposed (even if informally) for clusters hominin fossils than for those from other taxonomic groups. Nonetheless, there are few formal routes for recognizing this status based on quantitative evaluations of traits.

Although quantitative approaches to phylogenetics have typically ignored anagenesis, stratocladistic methods were developed to leverage temporal occurrence data to help reconstruct phylogenetic trees that have the potential to express direct ancestor-descendant relationships (Fisher 2008). Stratocladistics uses the criterion of maximum parsimony (MP) to minimize both the number of homoplasious character changes and unsampled stratigraphic intervals implied by topologies that include both bifurcating and serially linked segments of phyletic lineages. Several authors have expressed objections both to the use of temporal data in phylogenetic inference and the capability of available data to adequately test ancestral and collateral relationships between species (Smith 2000). Nevertheless, direct ancestors are expected to occur in the fossil record (Foote 1996), and the integration of temporal data has been shown to improve reconstruction accuracy over morphological analyses alone (Fox et al. 1999). In addition, those earlier criticisms, which have been frequently raised by proponents of cladistic methodologies, are also less relevant when placed in the context of the modern probabilistic approaches. While cladistics operates at the level of cladograms, both stratocladistics and recent probabilistic approaches reconstruct phylogenetic trees. This renders earlier criticism of temporal data largely irrelevant given the current landscape of phylogenetic methodology. As such, it stands to reason that probabilistic approaches are remiss when they fail to accommodate the possibility of ancestor-descendant relationships between temporally heterogeneous taxa.

There have been several attempts to explicitly extend the intent and logic of stratocladistics into probabilistic approaches using maximum-likelihood (ML) (Huelsenbeck and Rannala 1997; Wagner 1998). However, these have not found widespread use. Aspects of these have been appropriated more recently by methods that use some combination of morphological, fossil preservation, and lineage diversification models to infer relationships and lineage divergence times in a Bayesian context (Pyron 2011). This framework has been extended to explicitly accommodate ancestor-descendant relationships by modelling lineage diversification and fossil preservation processes (Stadler 2010; Bapst and Hopkins 2016; Stadler et al. 2017), in particular through the use of the ‘Fossilized Birth-Death’ (FBD) prior (Heath et al. 2014; Zhang et al. 2016). However, the assumptions and ramifications of doing so have not been well explored.

One outstanding question regarding these approaches is the extent to which morphological data themselves are able to illuminate questions of direct ancestor-descendant relationships without explicitly modelling the process of lineage diversification. This may be especially important in clades with gappy fossil records, such as many lineages of terrestrial vertebrates. In these cases, there may be less information available to reconstruct speciation and extinction parameters, and so a characterization of the identifiability of hypotheses of direct ancestry from morphology alone can benefit future studies in these taxa. In addition, methodological extensions of Bayesian tip-dating methods that accommodate serial sampling of lineages have been largely developed for use in epidemiological systems, where they are used to model molecular sequence evolution along single lineages. Although these systems are useful models of patterns in the fossil record, their sampling is often incomplete, and so they may sometimes call for different approaches in practice. Non-Bayesian *a posteriori* time-scaling (APT) approaches, such as *cal3*, that accommodate ancestor-descendant relationships also exist (Bapst 2013). These apply divergence times to unscaled cladograms that have been inferred from character data alone. Although both Bayesian and APT approaches have been shown effective when applied to fossil taxa (Bapst 2013; Bapst and Hopkins 2016; Gavryushkina et al. 2017; Stadler et al. 2017), there has been limited discussion of the statistical properties, limitations, and implications of using existing paleontological data and probabilistic models to identify anagenesis. Nevertheless, anagenesis is an evolutionary question of critical importance, and so the development of a probabilistic approach using likelihoodist criteria may be broadly beneficial by clearly quantifying the evidence for anagenesis present in paleontological data.

Although cladistic methods have been applied to improve understanding of certain aspects of hominin evolution (Delson et al. 1977; Chamberlain and Wood 1987; Strait et al. 1997; Irish et al. 2013), their exclusive focus on bifurcating relationships has precluded the study of direct ancestry except in cases where character polarity and specimen sampling have been carefully considered (Kimbel et al. 2006). More recently, Bayesian approaches have been suggested to provide a clearer view of hominin evolutionary relationships (Dembo et al. 2015; 2016), but their results still contain ambiguities. For example, like earlier cladistic approaches (Strait et al. 1997; Strait 1999), results from Bayesian analyses conflict with qualitative interpretations of hominin relationships in several key instances. This divide is exemplified by considering the taxa *Homo heidelbergensis* and *Homo antecessor,* for whom hypotheses of anagenesis are often considered. For example, previous studies have alternatively suggested that either *Homo heidelbergensis* or *Homo antecessor* may be directly ancestral to *Homo sapiens* and *Homo neanderthalensis* (Mounier et al. 2009), while some have debated the suggestion that *Homo antecessor* is ancestral to *Homo heidelbergensis* (Stringer 2012). Others have suggested alternately that *Homo heidelbergensis* is either a chronospecies directly ancestral to *H. neanderthalensis*, or directly ancestral to both *H. sapiens* and *H. neanderthalensis* (Rightmire 1998; Rosas and Bermúdez De Castro 1998; Stringer 2012). This uncertainty is underscored by the suggestion that cladistic methods and data are unreliable in their ability to describe hominin evolutionary patterns and history (Collard and Wood 2000).

In this study, we describe an approach to phylogenetic inference that combines models of stratigraphic preservation and morphological evolution to reconstruct time-scaled phylogenies and distinguish between anagenesis and cladogenesis using ML and the Akaike Information criterion (AIC). This approach seeks to simplify existing methods, such as the APT and FBD approaches described above, to clearly identify the signal for directly ancestral relationships presented by morphological data alone. Unlike more complex approaches, ours seeks only to identify information in morphological and temporal data that establishes differential support for ancestor-descendant and bifurcating relationships. As a result, our approach explicitly relies on models of morphological evolution and stratigraphic preservation rather than on models of lineage diversification.

We apply our new approach in hominins using a morphological supermatrix borrowed from the literature (Dembo et al. 2015; 2016). Hominins are a particularly compelling case study for our questions surrounding the identifiability of phylogenetic models that incorporate anagenesis. Although cladistic and phylogenetic methods have been applied to hominins (Chamberlain and Wood 1987; Strait et al. 1997; Strait 1999; Irish et al. 2013; Dembo et al. 2015; 2016), previous methods’ exclusive focus on bifurcating relationships has led to their inability to fully address outstanding hypotheses of hominin evolution, which are often predicated upon the occurrence of anagenesis. Our analysis identified several areas of anagenetic change in hominins, demonstrating cases where temporal data both corroborate and refute results achieved using morphology alone. We recognize that increasingly thorough compilation of morphological data and new fossil discoveries is likely to continue to revise and refine current understanding of hominin evolution. Nevertheless, our approach sheds light on the capabilities of existing models and data to accommodate anagenetic change without becoming entangled in a complex web of Bayesian assumptions. Moving forward, application of our method in hominins provides general demonstration of the importance of accommodating anagenetic change in phylogenetic methods to generate deeper understanding of evolutionary patterns in the fossil

## Materials and methods

### Inference of ML topology

Our approach evaluates the likelihood of candidate topologies using probabilistic models of fossil preservation and morphological evolution (Huelsenbeck and Rannala 1997; Lewis 2001). We perform a semi-automated tree search by calculating the likelihoods of these models on a set of candidate topologies. This approach tests hypotheses of direct ancestorship by combining branches with non-overlapping ranges and comparing cladogenetic and anagenetic models using the AIC. All code developed for these analyses is publicly available and implemented in the *mandos* package (www.github.com/carolinetomo/mandos).

### Identification of a fully-bifurcating ML tree

Our approach has yet to be implemented with a fully automated tree searching algorithm, so we combine semi-automated rearrangements with manual perturbations to search for the ML topology. This is done by exploring tree-space surrounding a starting tree estimated from morphological characters alone. In this study, we obtained a ML starting tree using RAxML, version 8.2.11, using the Mk model of morphological evolution (Lewis 2001). The morphological data were separated into partitions according to the number of possible states (i.e., binary, trinary, etc.) and analyzed under separate models. This partitioning scheme was maintained for all subsequent morphological likelihood calculations, including those used below in the AIC comparisons. We then performed a series of nearest neighbor interchange (NNI) and subtree pruning and regrafting (SPR) operations. These yielded a set of 1700 candidate topologies from which we identified the fully-bifurcating topology best supported by both morphologic and stratigraphic data. This tree provided a starting point drawn from both morphologic and stratigraphic lines of evidence. From here, we explored ancestor-descendant relationships using the model-testing approach described below.

### Modelling stratigraphic preservation

Stratigraphic likelihoods were calculated under a homogeneous Poisson process of geologic preservation. When applied to phylogenetics, inference under this model has been shown to accurately recover simulated phylogenetic relationships (Huelsenbeck and Rannala 1997). The likelihood function is derived from a Poisson process as the probability of observing the first occurrence (*o_f_*), last occurrence (*o_l_*) and number (*n_o_*) of occurrences in the stratigraphic record given some origination and extinction time (*t_f_* and *t_l_*, respectively) and preservation rate (λ). These likelihoods are calculated independently for each (*i*) of *b* total lineages and multiplied to yield the overall tree likelihood (see (Huelsenbeck and Rannala 1997) for full derivation):

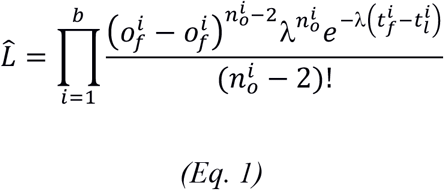

This equation reaches its maximum as branching and extinction times approach the first and last occurrences in the fossil record, and so the likelihood is maximized across the tree when the total amount of unsampled time implied by the topology is minimized. Although this approach differs from stratocladistic parsimony in its treatment of occurrence data as continuous rather than discrete, this property causes the preservation model to behave as a statistical formalization of the stratigraphic parsimony debt calculations undertaken in parsimony-based stratocladistic analyses. We estimate lineage origination and termination times along with preservation rate using multivariate numerical optimization routines implemented in SciPy (Jones et al. n.d.).

When combined with morphological data and models of character evolution, this approach represents a comprehensive extension of traditional stratocladistics based entirely upon probabilistic models.

### Identification of anagenesis

Using the bifurcating topology with the highest likelihood under both stratigraphic and morphologic models, we manually identified a set of potential ancestor-descendant relationships. To explore a more comprehensive range of both anagenetic and cladogenetic arrangements, we also extensively perturbed results manually and compared likelihoods. Although a fully-automated tree searching approach will ultimately be desirable in future versions of our method, our approach to tree-searching is similar to those used in previous stratocladistic studies. Starting with the fully bifurcating tree, we identified putative ancestor-descendant arrangements by collapsing each branch with a temporal range beginning earlier than the range represented by its sister lineage. We isolated each putative episode of anagenesis and compared the morphologic and stratigraphic likelihoods of anagenetic and cladogenetic arrangements using AIC scores. This was required because cladogenetic nodes assume one more parameter than anagenetic nodes (i.e., the branch length of the new lineage), and so bifurcating trees contain more parameters than anagenetic trees. Comparison of AIC scores enables a comparison of the relative quality of models with different numbers of parameters. AIC score is calculated from the number of model parameters (*k*), and the log-likelihood (*L*):

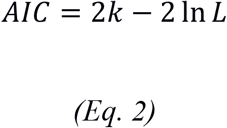

Since models with larger numbers of parameters are more prone to overfitting, they are biased toward possessing higher likelihoods than less complex models. As a result, likelihood scores of models that differ in dimension are not directly comparable. AIC scores represent the amount of information lost by a model when representing data, with lower scores indicative of models that preserve a greater amount of information. AIC accommodates for differences in parameter count between models by penalizing the addition of additional parameters, seeking to optimize the trade-off between improvements in model fit associated with added parameters and the loss of statistical power that results from overparameterization. In our use, AIC facilitates comparison of phylogenetic models of differing dimensions by penalizing the addition of branches that are better explained through an anagenetic pattern. The parameter count, *k*, for each phylogeny is calculated by summing the number of estimated branch lengths for each tree with the number of parameters used in the partitioned Mk substitution model.

Under the Poisson preservation model, stratigraphic likelihood predictably improves when unsampled time implied by the phylogeny is reduced, and so the acceptance of anagenetic arrangements also requires the support of morphology. This requires a novel calculation of morphological likelihood where the probability of transitioning from an observed, rather than an uncertain, parental character state to a single or multiple descendant character states is calculated under the Mk model (Fig. 1). This calculation differs from that used on multi-furcating nodes.

**Fig. 1.**
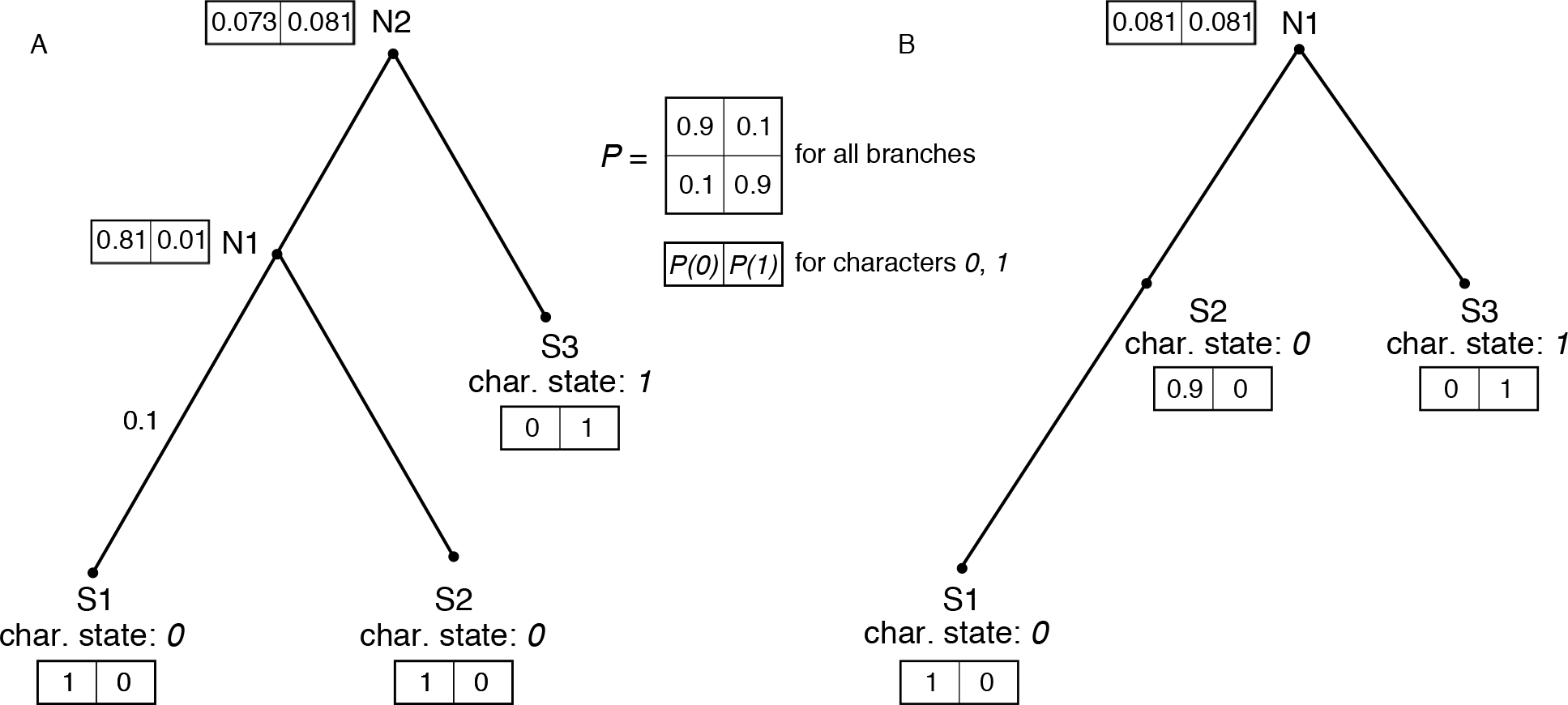
Comparison between anagenetic and cladogenetic trees. A) Likelihoods are calculated on bifurcating arrangements using the standard ‘pruning algorithm’. B) Anagenetic likelihoods are calculated using a novel approach, where ancestral states may be know.

Since the sequences at internal nodes representing unobserved taxa are unknown, the likelihood of character data at the tips is typically calculated by summing over all possible states at each unobserved internal node (Felsenstein 1981). However, when dealing with anagenesis, the sets of character states at some internal nodes are known. In these cases, the likelihood is calculated as the conditional probability of observing the set of traits at the tips given the set of traits possessed by the putative ancestor. Anagenetic arrangements are only accepted when there has been a sufficiently small amount of character change. This procedure resembles model testing procedures used to reconstruct ancestral DNA sequences, which compare conditional likelihoods of different permutations of character states at ancestral nodes (Yang et al. 1995). Like stratocladistics, this use of probabilistic models of character change enables morphology to occupy a central role in identifying anagenesis. As a result, this approach can in principle be used to test anagenesis even in the absence of a temporal model. A demonstration and test of this method using simulated data is provided in the supplement.

### Morphological matrix

We performed our analysis on a supermatrix of 391 discrete craniodental characters compiled by Dembo and colleagues (Dembo et al. 2015; 2016). We removed all ambiguous character states, as researchers did not identify whether these were truly ambiguous or polymorphic. While ambiguous character codings are unlikely to provide significant phylogenetic information, existing Markov models of discrete character evolution do not accommodate polymorphism. We excluded the taxa *Kenyanthropus platyops* and *Homo naledi* from the present analysis. The features that are diagnostic of *K. playtyops* have been suggested to result from taphonomic distortion resulting from matrix expansion, rather than from true biological differences (White 2003). Thus, we omitted this taxon in hopes of shedding greater light on the remaining, more widely accepted hominin taxa. *Homo naledi* was omitted because the data provided in the original study yielded an ML topology placing *H. naledi* as sister to *H. sapiens*. Although the phylogenetic affinity of *H. naledi* is a major outstanding question in paleoanthropology, the confusing signal presented by the *H. naledi* data, which are relatively recently acquired and therefore represent less well-studied fossils overall, reduced our confidence in the ability of this dataset to resolve its placement. Therefore, to avoid any confounding effects from the potential unreliability of the *H. naledi* data, we performed our analyses on the remaining subset of the data after *H. naledi* was removed. This enabled us to explore the phylogenetic relationships between better-known hominins.

### Geologic occurrence times

We surveyed the literature to obtain the observed temporal range of each taxon in continuous time. Reported radiometric dates for the oldest and youngest fossils were taken as the first and last observations. Some specimens are ambiguous in their taxonomic assignment; these were excluded from the analysis. We also gathered the number of total occurrences as the number of localities where each taxon has been identified as listed in MacLatchy et al. (2010), and supplemented these with additional localities identified in the literature. Cases where multiple specimens belonging to the same taxon have been identified at a single locality were treated as single occurrences. Although we recognize the potential ambiguity in delineating between sites, localities, and occurrences, we attempted to coarsely characterize the total number of occurrences using the number of sites at which each taxon occurs. This approach is more likely to underestimate the number of occurrences than overestimate them, which we expect to yield more conservative statistical support for competing topologies under the preservation model. A comprehensive list of the sites used to define temporal ranges for all taxa is provided in the supplement.

## Results and Discussion

### Anagenesis in the hominin fossil record

Our analysis yielded evidence for several instances of anagenesis in the hominin fossil record (Fig. 2). Our analysis reconstructed *Australopithecus anamensis* as directly ancestral to *Au. afarensis*. This result agrees with broad acceptance of *Au. anamensis* and *Au. afarensis* as phyletically linked chronospecies (Leakey et al. 1995; Ward et al. 2001; Kimbel et al. 2006). Although our analysis recovered *Ar. ramidus* as sister to the anagenetic *Au. anamesis-Au. afarensis* branch, the morphological data did not support the collapse of *Ar. ramidus*. Nevertheless, *Ar. ramidus* possessed poor character sampling in the matrix, and so its placement should be regarded as tentative. *Sahelanthropus tchadensis* is recovered as a direct ancestor to the rest of the hominin clade. This result should also be treated cautiously due to the small number of characters recovered for analysis of this portion of the phylogeny, but it is in line with with *Sahelanthropus’* status as the oldest recognized hominin (Brunet et al. 2002; Guy et al. 2005; Zollikofer et al. 2005).

**Fig. 2.**
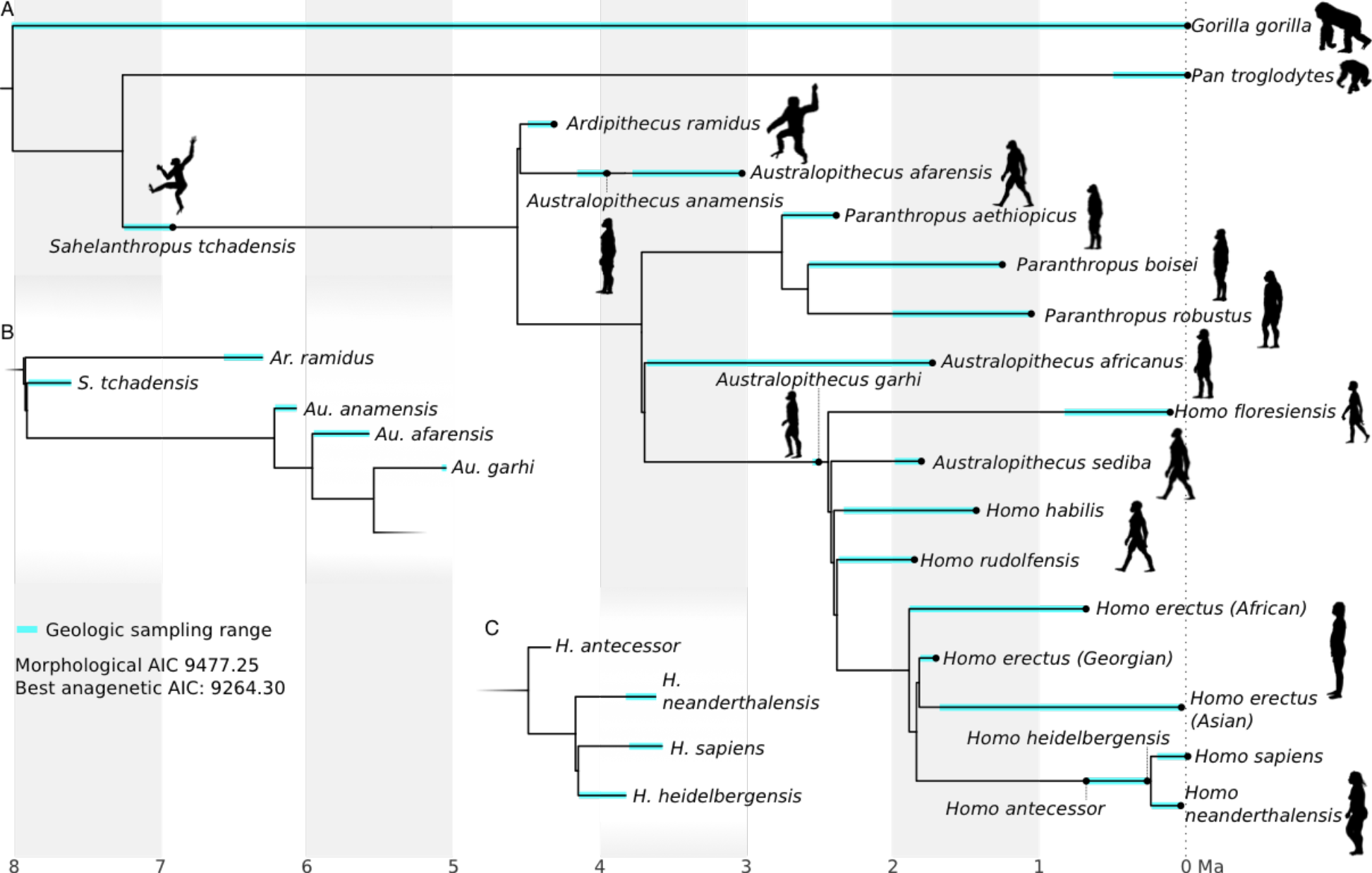
Phylogenetic relationships between hominin species. A) Reconstruction with the best AIC score when temporal data are considered alongside morphological data and anagenesis is accommodated. B, C) Areas of the tree inferred from morphology alone that differ from panel A. Both regions display misleading results when anagenesis is not considered. This is reflected in the improvement in AIC observed in the preferred anagenetic topology. Silhouettes obtained from phylopic.org.

Consistent with an early appraisal (Asfaw et al. 1999), our final analysis inferred *Au. garhi* to be directly ancestral to the *Homo* clade (Fig.2a). This conflicts with cladistic analyses that placed *Au. garhi* as outgroup to *Au. africanus*, *Paranthropus*, and *Homo* (Strait 1999). However, when anagenesis is not considered and phylogeny is inferred from morphology alone, we recover the same placement for *Au. garhi* as the cladistic result (Fig. 2b). Like the example above, this may reflect the constraint that strictly bifurcating methods impose on phylogenetic reconstructions among fossil taxa. However, preference for *Au. garhi* as ancestral to *Homo* is weak, with an only slightly improved AIC score over the *Au. garhi* outgroup hypothesis (Table 1).

**Table 1.** Support for alternative hypotheses regarding the placement of *Au. garhi*.

| Hypothesis | AIC score |
| --- | --- |
| *Au. garhi* outgroup to *Paranthropus* and *Homo* | 9271.11 |
| *Au. garhi* directly ancestral to *Homo* | 9264.30 |

Our results also better reconcile quantitative and qualitative interpretations of the evolutionary mode and relationships among the *Homo* species leading to modern humans and Neanderthals. Paleoanthropologists have variously interpreted *Homo heidelbergensis* as either: 1) a direct ancestor to both Neanderthals and modern humans (Rightmire 1998; Mounier et al. 2009; Stringer 2012; Buck and Stringer 2014), or 2) a chronospecies leading to Neanderthals (Rosas and Bermúdez De Castro 1998). However, quantitative phylogenetic analysis has placed *H. heidelbergensis* as sister to Neanderthals (Dembo et al. 2015). Our ML result based on morphological characters alone places *H. heidelbergensis* as sister to *H. sapiens*. However, when temporal data are incorporated and anagenesis is accommodated, AIC support improves substantially, and *H. heidelbergensis* is collapsed to represent a direct ancestor of modern humans and Neanderthals (Fig. 3). We also uncover one other instance of anagenesis in this clade. Opinions are divided as to whether *H. antecessor* represents a direct ancestor of later hominin species or is an evolutionary dead end (Bermúdez de Castro et al. 1997; Stringer 2012; Dembo et al. 2015), but our results provide support for combining *H. antecessor* and *H. heidelbergensis* into a single lineage. This suggests a long episode of anagenetic evolution immediately prior to the divergence between modern humans and Neanderthals.

**Fig. 3.**
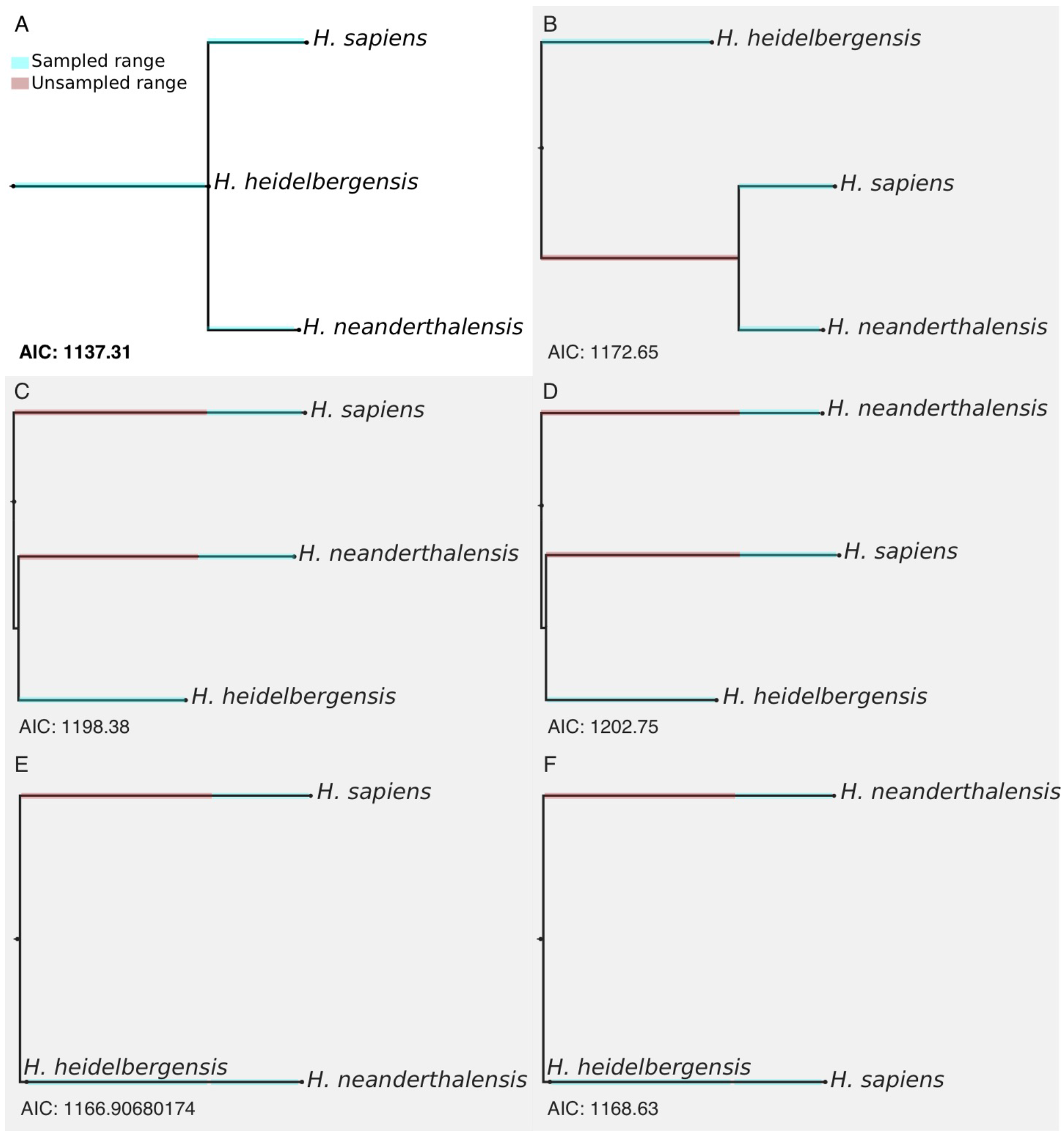
AIC scores calculated for each possible arrangement between *H. sapiens*, *H. neanderthalensis,* and *H. heidelbergensis*.

Overall, our reconstruction of relationships among later species of *Homo* immediately preceding and encompassing modern humans and Neanderthals supports the hypothesis that *H. heidelbergensis* is directly ancestral to both modern humans and Neanderthals. This result differs from the hypothesis supported by the analysis performed by Dembo and colleagues (Dembo et al. 2015), and is instead more consistent with an earlier exploratory statistical analysis (Mounier et al. 2009), and with the position frequently suggested by paleoanthropologists (Rightmire 1998; Stringer 2012; Buck and Stringer 2014). While previous phylogenetic analyses have yielded results that equivocate or disagree with the common interpretation of *H. heidelbergensis* as the last common ancestor of modern humans and Neanderthals, our analysis shows that the consideration of direct ancestry can generate statistical support for phylogenetic results that conform more closely to positions generated through researchers’ subjective interpretations and exploratory data analyses. This finding supports a general argument against the use of cladistic and phylogenetic methods that are restricted to bifurcating relationships in fossil taxa, where the possibility of variability in evolutionary mode (i.e., occurrence of both anagenesis and cladogenesis) is at odds with an assumption that evolution proceeds by cladogenesis alone (Fig. 2).

Our results differ markedly from previous phylogenetic studies seeking to reconstruct hominin phylogeny using probabilistic and cladistic methods. In key regions of the tree, results achieved under our method reveal support for hypotheses more consistent with many qualitative interpretations of hominin relationships, demonstrating the importance of explicitly accommodating anagenesis in the phylogenetic reconstruction of fossil taxa. This may explain some of the historical difficulty in reconciling paleontological interpretations of hominin relationships with cladistic results. For instance, Dembo and colleagues’ results are inconsistent with earlier suggestions that *Au. anamensis* and *Au. afarensis* are chronospecies differentiated through anagenesis. However, by considering ancestor-descendant relationships and incorporating temporal data, our analysis reveals that a linked *Au. anamensis-Au. afarensis* lineage is the arrangement most strongly supported by the data. Generally, this suggests that cladograms and strictly bifurcating phylogenies may be inadequate when describing evolutionary relationships between fossil taxa, and so ancestor-descendant relationships should be considered during topological inference. Further, we argue that explicit testing of ancestor-descendant relationships is important even in cases where bifurcating trees are not wholly misleading, as their omission precludes us from considering the full range of possible evolutionary scenarios.

Although previous studies addressing hominin phylogeny using probabilistic methods represent a significant step forward in weighing alternative evolutionary hypotheses, we suggest that their phylogenetic reconstructions have suffered from methodological limitations that were not generally perceived. We suggest that overcoming these limitations can provide a substantial step forward in closing the gap between the paleobiologists’ interpretations and previous cladistic and phylogenetic results. In particular, we show through our analyses that the apparent discordance between quantitative and qualitative assessments of evolutionary relationships can be reconciled by extending phylogenetic models to explicitly accommodate anagenesis.

### Ancestors, anagenesis, and evolutionary processes

Our method does not seek to distinguish between speciation modes at a mechanistic level. As noted by Fisher (2008), overlap between the extinction and origination times of taxon pairs does not necessarily preclude an ancestor-descendant relationship. Instead, it is possible for ancestral and descendant lineages to coexist.

This reality may complicate the identification and evolutionary interpretation of ancestor-descendant relationships from temporal data alone. Doing so requires diversification and preservation models that contain additional parameters that quantify completeness of the fossil record. Such models are currently implemented in the *cal3* time-scaling method, which seeks to distinguish between splitting, budding, and anagenesis from a cladogram with a fixed topology (Bapst and Hopkins 2016). Our approach is distinct both theoretically and operationally from *cal3*, instead relying most heavily on morphological evidence to weigh the likelihood of ancestor-descendant relationships without considering the completeness of fossil sampling. At the population level, one might expect some temporal overlap between ancestral and descendant taxa undergoing anagenetic change, especially if the participants are widely distributed geographically. As a result, even if the temporal ranges corresponding to taxa identified as ancestor-descendant pairs are discovered to slightly underestimate the time of coexistence, the inference may simply highlight the fuzziness in the taxonomic placement of fossils belonging to lineages undergoing continuous transformation and in discerning between anagenesis and evolutionary budding given incomplete sampling (Fig. 4). This interpretation is consistent with previous authors’ treatment of temporal ranges when identifying anagenesis between taxa, which have allowed a period of overlap between putative ancestor-descendant pairs (Aze et al. 2011; Strotz and Allen 2013).

**Fig. 4.**
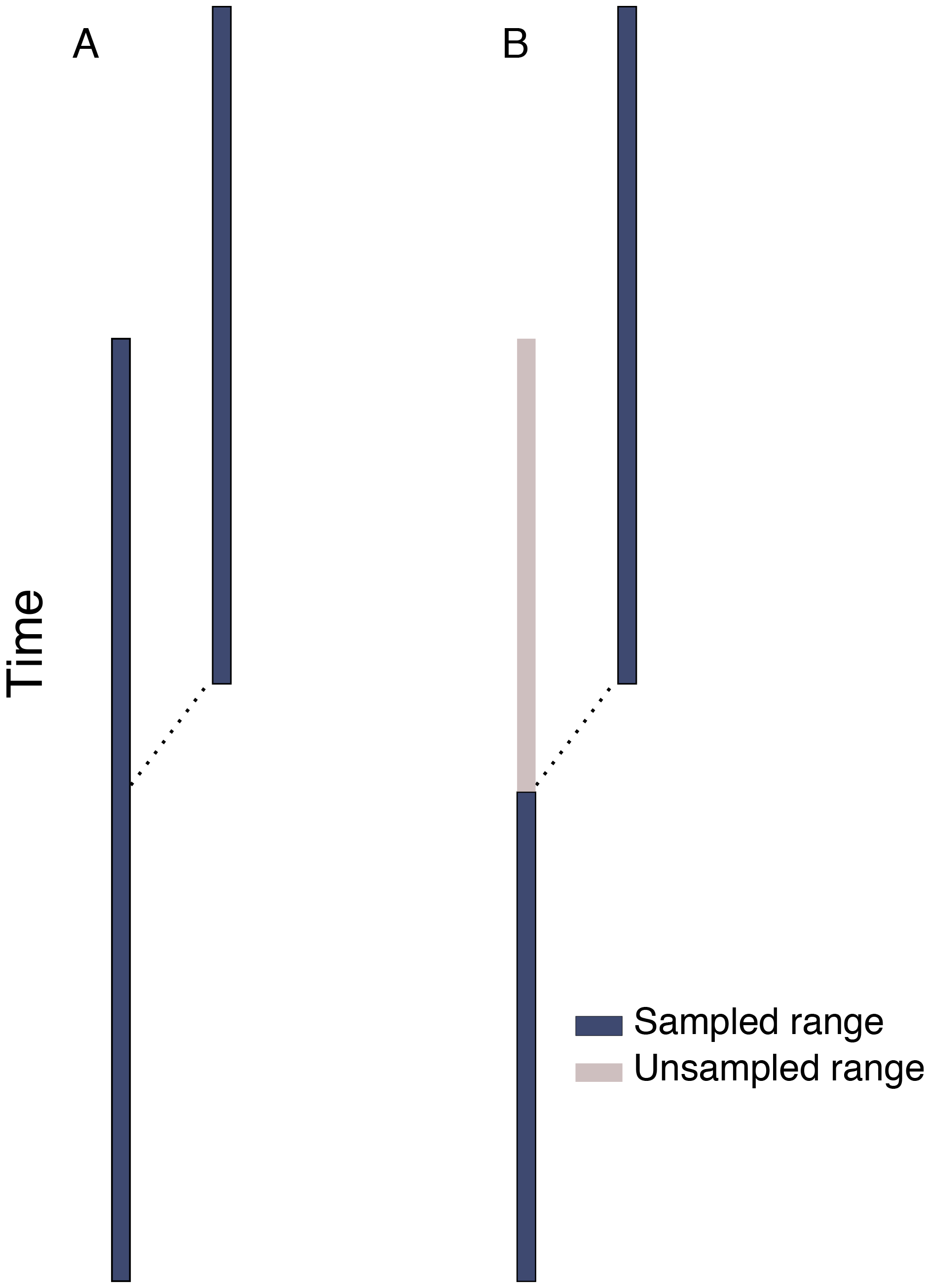
A) Speciation mode interpreted as budding when sampling is complete. B) Incomplete stratigraphic sampling may create an inability to distinguish between anagenesis and budding when sampling is sparse.

We use the term ‘ancestor’ modelled loosely after Gingerich (1979). Ancestors identified through our method represent collections of samples possessing a suite of morphological character states that is not sufficiently differentiated from a single or a set of subsequently occurring samples to warrant assignment to distinct lineages. In our usage, we consider anagenesis as any evolutionary change occurring along these serially linked phyletic lineage segments. Thus, the taxonomic and phylogenetic units of analysis are fundamentally important to the formulation and interpretation of our results. Hominin taxa are often represented by only one or a small number of samples, and the number of species may be overestimated (White 2003).

This may complicate the ability to meaningfully characterize phylogenetic relationships from a precise mechanistic view. Nevertheless, morphological and temporal data can be combined to summarize the relative support for anagenetic and cladogenetic patterns in the inheritance of observed character states (i.e. even in cases where distinctiveness may not be at the species level).

Under our method, ancestor-descendant relationships might be interpreted either as true anagenesis (i.e., a single population undergoing gradual transformation), or as some form of budding cladogenesis. Previous researchers have argued that true anagenesis is rare compared to budding when analyzing the fossil records of densely-sampled marine invertebrate lineages using more complex preservation models (Bapst and Hopkins 2016). Nevertheless, we suggest that distinction between these two modes may often be impossible in terrestrial vertebrate lineages with large sampling gaps. For example, our results among early hominin species include multiple inferred direct ancestors, but the large gaps in stratigraphic sampling throughout this region of the tree hamper the ability to determine whether these relationships represent true anagenesis or budding that has been obfuscated by poor sampling.

Instead of relying upon hope that more complex preservation models are robust to wide sampling gaps, our method disregards the completeness of sampling to evaluate the support for ancestor-descendant pairs using morphology. Although temporal data occupy an important place in our approach, they are largely used as a guide to constrain the set of possible ancestors and descendants and to provide additional insight when morphological data are equivocal. Coarsely speaking, our approach focuses on patterns in morphological differentiation and lineage *disparification*, while approaches such as *cal3* model lineage *diversification*. Further empirical and simulation-based work is needed to determine the relative merits of methods of testing ancestor-descendant relationships, such as *cal3*, that explicitly focus on diversification patterns, and ours, which weighs morphological patterns more heavily. We speculate that their relative accuracy may depend largely on the completeness of sampling in the rock record and the correlation strength between morphological change and lineage diversification, although other factors may also be important.

The scales at which phylogenetic data are sampled may further complicate mechanistic evolutionary interpretations. Morphological character matrices often lack samples across the entire stratigraphic range of each taxon, and so generally assume morphological stasis within lineages. It is therefore often impossible to observe gradual morphological change within and between taxa. These considerations might cause anagenetic relationships identified here to represent either true anagenesis or some form of ‘pseudo-anagenesis’, where stratigraphic and morphological data appear consistent with anagenesis but the persistence of the ancestor has not been sampled. The ancestor-descendant relationships identified by our method may be interpreted in several ways. As sampled, these results may be roughly conceived as anagenesis in the sense that the mode of evolutionary change between taxa is indistinguishable from evolution occurring along a single lineage, depending upon the completeness of sampling and the degree to which morphological disparity correlates with true biological species diversity. This interpretation is consistent with historical usage by paleobiologists (Gingerich 1979; Smith 1994; Levinton 2001). Thus, our approach seeks to reveal the extent to which existing cladistic and temporal data can provide evidence for non-branching evolutionary modes and does not seek to resolve conceptual issues that may stem from incomplete sampling, lineage diversification, or population-level evolutionary change. Regardless of the fine-scale evolutionary interpretation, failure to accommodate phyletic change and ancestor-descendant relationships when inferring phylogenetic relationships can generate views of evolutionary history that are positively misleading in the sense that incorrect results cannot improve with the addition of new data.

### Some practical methodological considerations

Concerns regarding the accuracy of probabilistic approaches have been raised, stemming from the reliance of these methods on the overly simplistic Lewis Mk model of morphological evolution (Goloboff et al. 2017). These critics advocate the use of cladistic methods, arguing that Markov models inadequately capture the complexities of morphological evolution. Although we agree that existing substitution models oversimplify these processes, our results suggest that the accommodation of ancestor-descendant hypotheses in probabilistic methods can improve the fidelity of phylogenetic reconstructions, even when Lewis Mk is used as the underlying model of morphological change. As a result, concerns regarding the adequacy of existing morphological substitution models may be partially alleviated by considering hypotheses of direct ancestorship. This is supported by simulation work showing that stratocladisics outperforms cladistics in topology reconstruction (Fox et al. 1999). Further exploration is needed to demonstrate more thoroughly the limitations of our new approach, which builds upon stratocladistics by incorporating the benefits of probabilistic analyses, including 1) more explicit statements of the assumptions involved, and 2) the ability to weigh competing models using modern inferential criteria.

The method we describe seeks to enhance understanding of the fossil record by explicitly testing support for existing hypotheses of direct ancestorship while attempting to make more explicit assumptions than stratocladistics or recently developed Bayesian methods. Although it shares features with recent methods that use mechanistic evolutionary models in a Bayesian context to infer ancestral relationships (Zhang et al. 2016), our approach is intended as a foundational, minimally complex framework for exploring the behavior of probabilistic models to evaluate support for anagenesis in temporal and morphological data. As such, our method should be viewed as a complement to, rather than a simplification of, existing Bayesian approaches. That is, our method encourages examination of the informativeness of the data without the increased complexity of assessing prior probabilities. Our method further differs from both existing Bayesian and parametric APT approaches by explicitly omitting diversification parameters and instead placing morphological data in a central role when evaluating hypotheses of direct ancestry. In doing so, temporal data help to delineate the set of possible ancestors and play an important role in measuring the fit of candidate trees to the observed stratigraphic record. Our method does not seek to reconstruct diversification processes, and instead focuses on identifying hypotheses that best describe only the information contained within morphological and temporal datasets. Assessment of information contained within datasets and tests of hypotheses can also be achieved using Bayesian approaches (Lewis et al. 2016), but likelihoodist approaches such as ours streamline these procedures by reducing complications presented by prior probabilities and Bayesian Markov-chain Monte Carlo sampling.

Although Bayesian methods can be beneficial in certain circumstances, our method simplifies identification of anagenetic hypotheses using evolutionary and stratigraphic models. We observe that the likelihood surface surrounding certain nodes may possess low peaks (Table 1), which likely results from sparse sampling and relatively low information in the morphological characters. Since Bayesian approaches often average results across this surface, it is possible that they may fail to capture those relationships best supported by the data by including information from weakly supported hypotheses. This is of greater concern in paleontological than neontological data because the increased abundance of molecular data is often likely to result in more clearly defined peaks in the likelihood surface. In cases where information is sparse, likelihoodist approaches such as ours offer the benefit of filtering through noisy and equivocal signal to reveal the hypothesis best supported by the data. Although these benefits are also achievable through Bayesian approaches, additional caution must be taken to select priors that do not dominate weakly informative data. In addition, careful thought should be given when summarizing the posterior/likelihood surface. Averaging across a relatively flat surface might yield poor results (Yang and Zhu 2018), while the comparison of individual point estimates, as is done here, may more clearly shed light on models best supported by the data while clearly capturing their relative support.

### How can we proceed?

Anagenesis and splitting cladogenesis were most pertinent to our analysis of hominin evolution, and the straightforward dichotomy between these simplified the assumptions and interpretation of our tests. Our approach may need to explicitly accommodate budding before being applied to groups with very dense fossil records. However, the assumptions required, which may often include morphological stasis within lineages, may make application to some phylogenetic datasets impractical. This is especially true in cases such as hominins, for which the fossil record implies large sampling gaps, and characters representative across stratigraphic ranges are often sampled from only a single individual. Expansions of our approach through implementation of new models will further test the implications of existing paleontological datasets for reconstructing complex evolutionary and geologic processes over deep timescales. We hope that the example provided here will encourage integration of more diverse evolutionary modes into phylogenetic methods yielding better explanations of temporal patterns in critical parts of life’s history.

As we emphasize above, the approach described here should be viewed as an attempt to explore the capability of phylogenetic methods to identify the signal of anagenesis using existing models to interrogate morphological and stratigraphic data. In doing so, we acknowledge that there are many complicated biological and geological factors that could be incorporated into this framework. For instance, previous researchers have accommodated heterogeneity in fossil preservation rates across time and among lineages (Foote 1997; 2001; Gavryushkina et al. 2014). There have also been several concerns raised in the literature regarding the adequacy of existing models of discrete trait evolution to inform complex evolutionary scenarios (Goloboff et al. 2017; Brown et al. 2017). Alternative models that use continuous characters may help to improve some of these issues (Felsenstein 1988; Goloboff et al. 2006; Parins-Fukuchi 2017). Moving forward, elaborations making use of new data sources and models will only continue to improve resolution of evolutionary patterns in the fossil record.

Finally, we acknowledge that our empirical results beg qualification. In particular, we expect that future studies will generate a more comprehensive and authoritative view of hominin evolution as improved data continue to become available. For instance, although we are currently cautious about making strong statements concerning the ancestral position of *Sahelanthropus* using this dataset, additional information may resolve this issue. This may be the case for several other areas of the hominin tree, which may be better resolved as temporal and taxonomic gaps in sampling are better filled by new discoveries. In addition, we concede the possibility that more comprehensive automated tree searching routines may reveal support for hypotheses that we failed to consider under our semi-automated approach. Therefore, instead of providing an authoritative view of hominin evolution, our study provides a springboard for future studies by showing that the accommodation of anagenesis can improve our view of the processes and relationships underpinning the evolution of fossil taxa. Nevertheless, due to the improved support and congruence of hypotheses that explicitly consider ancestor-descendant relationships, we recommend that future phylogenetic studies in hominins avoid methods that only consider bifurcating relationships. Future studies that build upon existing work in other taxa will also be important to better characterize the extent to which this suggestion can be generalized across the tree of life. Although the accommodation of directly ancestral relationships is especially relevant in hominin taxa, for which hypotheses of anagenesis have been long entertained through qualitative anatomical assessment, these results may also be important in other taxa. Further empirical work will be needed to develop a better understanding of the extent to which the consideration of direct ancestors can improve resolution of evolutionary patterns throughout the fossil record.

Author contributions
CPF designed and implemented the ML stratocladistic procedure, performed analyses, interpreted results, and drafted the manuscript. DCF made conceptual contributions to the stratocladistic approach developed here, contributed to the manuscript, and helped interpret results. EG gathered temporal occurrence ranges, helped interpret results, and edited the manuscript. LMM helped focus the scope of the hominin analysis, helped interpret results, and contributed to the manuscript.

## Acknowledgments

CPF would like to thank JW Brown, CW Dick, M Friedman, SA Smith, and GW Stull for helpful discussion of the manuscript and method.

